# Ribosome profiling in *Mycobacterium tuberculosis* reveals robust leaderless translation

**DOI:** 10.1101/2020.04.22.055855

**Authors:** Elizabeth B. Sawyer, Jody E. Phelan, Taane G. Clark, Teresa Cortes

## Abstract

*Mycobacterium tuberculosis*, which causes tuberculosis, expresses a large proportion of leaderless transcripts lacking the canonical bacterial translation initiation signals. The role leaderless genes play in the physiology of this pathogen, which can undergo prolonged periods of non-replicating persistence in the host, is currently unknown. We have previously demonstrated that levels of leaderless transcription increase under conditions of nutrient starvation. However, little is known about the implications of this for persistent infection. Here, we performed ribosome profiling to characterise the translational landscape of *M. tuberculosis in vitro*. Our data reveals robust leaderless translation in the pathogen and points towards different mechanisms for their initiation of translation compared to canonical Shine-Dalgarno genes. Furthermore, under conditions of nutrient starvation, we found a significant global up-regulation of leaderless genes in the translatome. Our data represents a rich resource for others seeking to understand translational regulation not only in *M. tuberculosis* but in bacterial pathogens.

## Introduction

*Mycobacterium tuberculosis* is an important human pathogen with a complex lifecycle characterised by long periods of persistence, in which bacteria enter a non-replicative, antibiotic-tolerant state (Stewart et al., 2003; Gomez and McKinney, 2004;). The mechanisms underlying bacterial adaptation to diverse environmental conditions encountered during infection, such as the acidification of phagosomes, the presence of reactive nitrogen and oxygen species, hypoxia and nutrient starvation are still poorly understood (Rengarajan et al., 2005). Recent findings have demonstrated differences in translational control between *M. tuberculosis* and the model organism *Escherichia coli*, and these mechanisms may play a role in the ability of *M. tuberculosis* to adapt to diverse environmental conditions (Trauner et al., 2012; Xue and Barna, 2012; Byrgazov et al., 2013; Bunker et al., 2015a) (reviewed in (Sawyer et al., 2018)).

The translation of messenger RNA (mRNA) into protein is a highly controlled process involving multiple layers of regulation at each step of, initiation, elongation, termination and ribosome recycling. In recent years, the heterogeneity of these regulatory mechanisms has begun to be appreciated and explored. In particular there is increasing evidence to show that distinct populations of ribosomes preferentially translate mRNAs with certain features (Xue and Barna, 2012; Byrgazov et al., 2013; Shi et al., 2017;). Untranslated regions (UTRs) of mRNAs can influence the stability and translational efficiency of the transcript either by providing binding sites for regulatory proteins or RNAs, adopting secondary structures such as internal ribosome entry sites or by repressing translation (Hughes, 2006; Morris and Geballe, 2000; Nguyen et al., 2020). Furthermore, short open reading frames (ORFs) within 5’ UTRs may inhibit translation by occluding the start codon or may themselves be translated (Ruan et al., 1994; Cao and Geballe, 1995; Werner et al., 1987).

The development of ribosome profiling techniques has proven to be a powerful technique for studying translation (Ingolia et al., 2009). It involves isolation of ribosomes bound to mRNA, enzymatic digestion of mRNA unprotected by the ribosome and sequencing the ribosome-protected fragments (RPF or footprints) of the mRNA. This technique enables very precise identification of specific mRNA strands being actively translated at any given moment in time. Ribosome profiling experiments performed in the non-pathogenic *Mycobacterium smegmatis* have uncovered important new details about translation, including the translation of hundreds of small, previously unannotated proteins at the 5’ ends of transcripts (Shell et al., 2015), the translation by alternative ribosomes that have distinct functionality compared with their canonical counterparts (Chen et al., 2019) as well as cysteine-responsive attenuation of translation (Canestrari et al., 2020).

The adaptive responses of mycobacteria to stress have been extensively analysed by transcriptional profiling in well-defined experimental models (Stewart et al., 2002; Betts et al., 2002; Deb et al., 2009; Rohde et al., 2012; Galagan et al., 2013; Rustad et al., 2014; Namouchi et al., 2016; Aguilar-Ayala et al., 2017; Martini et al., 2019). However, due to the well-established poor correlation between levels of mRNA transcripts and proteins in the cell (Cortes et al., 2017; Haider and Pal, 2013) and differences in mRNA and protein half-lives (Nguyen et al., 2020), it is unclear how the changes that occur in the transcriptome of bacteria that enter a persistent state play out in the proteome. Clearly, there are post-transcriptional control mechanisms that influence protein abundance, but these are currently poorly understood in mycobacteria. Previous genome-wide mapping of transcriptional start sites (TSSs) in *M. tuberculosis* has shown that this pathogen expresses a large number of transcripts that lack a 5’UTR (referred to as leaderless transcripts) including the sequence elements that usually play a role in ribosome recruitment and binding, such as the Shine-Dalgarno sequence (Cortes et al., 2013; Shell et al., 2015). It is known that under conditions of nutrient starvation, transcription of leaderless genes is upregulated and associated with genes expressed in nondividing cells (Cortes et al., 2013), but understanding of the impact of this regulation on translation is lacking. Hence, further understanding of the quantitative correlations between transcription and translation, and the regulation of translation of leaderless mRNAs is needed in order to gain more insights into persistent infection with *M. tuberculosis*.

In this Tools and Resources article, we have applied parallel RNA sequencing and ribosome profiling in *M. tuberculosis* H37Rv (reference strain) to gain further insights into the translational landscape of this pathogen and provide comprehensive measurements of translation under exponential growth and following a nutrient starvation model. Our data provide a rich resource for studying translation and its regulation in *M. tuberculosis*.

## Results

### The translational landscape of M. tuberculosis during exponential growth

To better understand genome-wide translation in *M. tuberculosis*, we adapted a previously published protocol for ribosome profiling (Latif et al., 2015) to be suited for a Biosafety Level 3 laboratory environment. Cultures of *M. tuberculosis* H37Rv were grown to an OD_600_ of ~ 0.6 and triplicate samples were processed for parallel ribosome profiling (Ribo-seq) and RNA sequencing (RNA-seq) (see Methods). Chloramphenicol was used to halt elongation for the generation of Ribo-seq libraries (Becker et al., 2013). The libraries were multiplex sequenced on an Illumina NextSeq 500 by Cambridge Genomic Services. On average, we sequenced 36 million fragments, with 30% of the RPF reads aligning to non-ribosomal RNA genome locations (Supplementary table S1). As our experimental procedure did not include a size selection step, we obtained a broad distribution of RPF read lengths, ranging from 20-37 nucleotides (Supplementary figure S1A). To evaluate if the range of RPF reads obtained were representative of translating ribosomes, we assessed the ribosome occupancy for genes aligned at their start codon for each read length individually. We assigned ribosome densities to the 3’ end of reads as it has been previously shown to yield precise information about ribosome position in bacteria (Balakrishnan et al., 2014; Nakahigashi et al., 2014; Woolstenhulme et al., 2015; Mohammad et al., 2019). This analysis revealed a defined peak near the start codon corresponding to initiating ribosomes for RPFs ranging from 29-37nt, whereas the shorter reads (20-28 nt) did not show a clear peak at the start codon and were considered to be artefacts of the experimental procedure (pre-initiation complexes and other material that precipitated with translating ribosomes) (Supplementary figure S1B). Therefore, for all downstream analyses only RPFs of length 29-37 nucleotides were considered. Samples were depleted of ribosomal RNA using the ribo-zero kit (see Methods) and remaining ribosomal RNA reads were filtered out of the dataset *in silico*.

To assess the quality and reproducibility of our data, we calculated the pairwise Spearman correlation for each pair of biological replicates for both the RNA-seq and Ribo-seq libraries, which showed a high degree of reproducibility between replicates (Spearman’s *r*> 0.93; Figure 1A). Next, we calculated the fraction of reads corresponding to various genomic regions for both the RNA and RPF reads. As RPF reads reflect translating ribosomes, it is expected to have a higher fraction of the reads mapped within or proximal to coding sequences and 5’UTRs, whereas RNA reads will have a broader distribution with a higher fraction of reads mapped to non-coding regions of the genome. As expected, whereas only 37% of the RNA reads accounted for coding sequences (CDS), 66% of the RPF reads mapped within CDS with a further 18% mapping within 5’UTRs (Figure 1B). Mapping of RPF reads within 5’UTRs is expected to reflect ribosomal engagement near the annotated start codon for translation initiation. To verify this, we further looked at the 76 expressed 5’UTRs shared across the top 100 most expressed 5’UTRs among the three biological replicates, which accounted for 72.3% of the total fraction of reads mapped to 5’UTRs. In 65 out of the 76 cases, mapped RPF reads to these regions could be attributed to ribosomal recruitment for translation initiation (Supplementary figure S2).

The overall expression rate for each gene during exponential growth was calculated as read density across each gene in reads per kilobase per million (RPKM). Plotting of gene expression rates for RPF and RNA across replicates revealed a high correlation between measures of transcription and translation consistent with the coupling of these processes in bacteria (Spearman’s *r* = 0.91; Figure 1C). In agreement with previous measurements of global transcription during exponential growth (Cortes et al., 2013), we were able to reliably quantify translation rates for 2,598 genes, representing 64.4% of the annotated translatome (see Methods, Supplementary Table 2, Supplementary figure S3).

To further investigate the extent to which the RPF read densities reflect the relative abundance of proteins in the cell, we looked at members of multiprotein complexes with known stoichiometry. The *rplJ* operon encodes the 50S ribosomal proteins uL10 (RplJ, Rv0651) and bL7/12 (RplL, Rv0652) which are present in one and four copies, respectively, in the assembled 50S subunit (Gordiyenko et al., 2010). The ATP synthase operon also encodes proteins that are present in different stoichiometries in the assembled complex (AtpA-H) (Foster and Fillingame, 2009; Lu et al., 2014). We plotted the stoichiometries of these proteins in the complexes against the RPF read densities and observed a close relationship between the number of read counts and the protein abundance (Figures 1D and 1E) providing further confirmation that our ribosome profiling data reflect genuine translation events.

**Figure 1.**
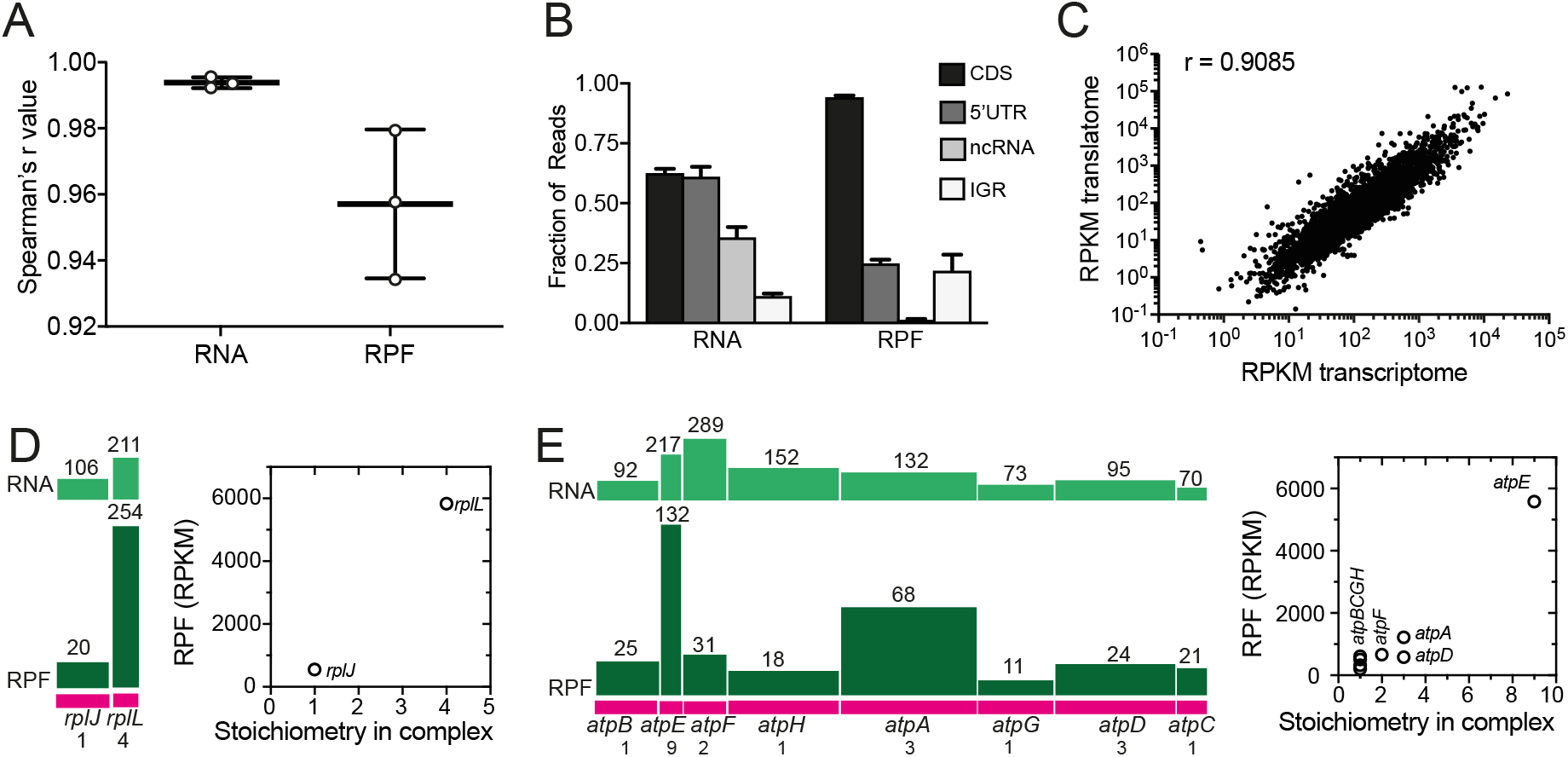
Measures of quality and reproducibility of the data. (A) Pairwise correlation coefficients calculated between each pair of replicates for total RNA and RPFs based on number of counts per gene. (B) Fraction of RNA and RPF reads aligning to various genomic features, coding sequences (CDSs), 5’ UTRs, non-coding RNAs (ncRNA) and intergenic regions (IGR). (C) Correlation between transcription and translation assessed from plotting average RPKMs across the three replicates for mRNA and RPFs. (D and E) Relationship between stoichiometries of proteins in complexes (50S ribosomal subunit and ATP synthase) and RPF read counts (in RPKM). The operon structure is shown in pink, where the length of the bar represents the gene length and the gene name and stoichiometry in the complex shown below. Above are shown the counts of RPF reads (dark green) and RNA reads (light green) where the height of the bar represents the average number of counts across the gene, indicated above. The graphs show the stoichiometry of the protein in the complex plotted against the RPF read counts (in RPKM).

In summary, we have sequenced total RNA and ribosome footprints from exponentially growing cultures of *M. tuberculosis* H37Rv. After filtering the data to exclude probable artefacts, analysis of the sequencing data indicates that our RPF data likely reflects genuine translation events in the mycobacterial cell, including proportional synthesis of genes from the same operons whose stoichiometries in the resulting protein complexes differ.

### Broad leaderless translation during exponential growth

We have previously demonstrated that a substantial subset of *M. tuberculosis* mRNAs is transcribed as leaderless transcripts that lack a 5’UTR and associated ribosomal recognition signals, but quantification of translation rates for this pathogen at a genome-wide level was lacking. From the data presented here, we were able to quantify similar reliable rates of translation for leaderless mRNAs (436 out of 2,598; 17%) and for Shine-Dalgarno mRNAs (486 out of 2,598; 19%); thereby corroborating that *M. tuberculosis* is able to efficiently initiate global translation of leaderless transcripts during exponential growth even in the absence of a Shine-Dalgarno sequence (Shell et al., 2015; Nguyen et al., 2020). Translation efficiencies (TEs) were calculated by dividing the RPKM of the translatome by the RPKM of the transcriptome and revealed significantly higher median efficiency levels for Shine-Dalgarno genes (Mann-Whitney test, P = 0.0195; Figure 2A), confirming previous observations between these genes and higher levels of expression (Ma et al., 2002).

We analysed the TE values of genes from various functional categories (Figure 2B upper panel) and found an association between the TE value and the gene functional categories that are required for optimal growth. In particular, energy metabolism (b) and ribosomal protein synthesis (k) categories had higher median TE values than others such as non-ribosomal peptide synthesis (h) (Kruskal-Wallis test, P <0.003). We also plotted the proportions of genes in each category that have a Shine-Dalgarno sequence (blue), are leaderless (orange) or have a 5’ UTR that does not include such a sequence (grey; Figure 2B lower panel). Looking together at Figures 2C and 2D it is apparent that those functional categories involved in primary metabolic functions tend to have higher proportions of Shine-Dalgarno genes and be more efficiently translated than categories with more leaderless genes. The exception is the toxin-antitoxin systems category (r) where the TE values are dominated by a small number of antitoxins with high efficiencies in exponential growth.

As TE is an estimate of how much protein is made from a single mRNA, comparing gene TE values against mRNA gene expression rates should provide information on the regulation of gene expression at the level of translation and reveal potential differences in TEs amongst gene categories. Plotting of mRNA to TE ratios for the different gene categories provided no evidence for an overall difference in the translation of leaderless and Shine-Dalgarno transcripts (Spearman’s *r* < 0.27) (Figure 2C). Overall, our data suggest that in contrast to *E. coli*, where the efficiency of leaderless translation is strongly dependent on the availability of initiation factors (Moll and Bläsi, 2002), *M. tuberculosis* can efficiently initiate and robustly translate leaderless transcripts using the translational machinery available during *in vitro* growth conditions.

**Figure 2.**
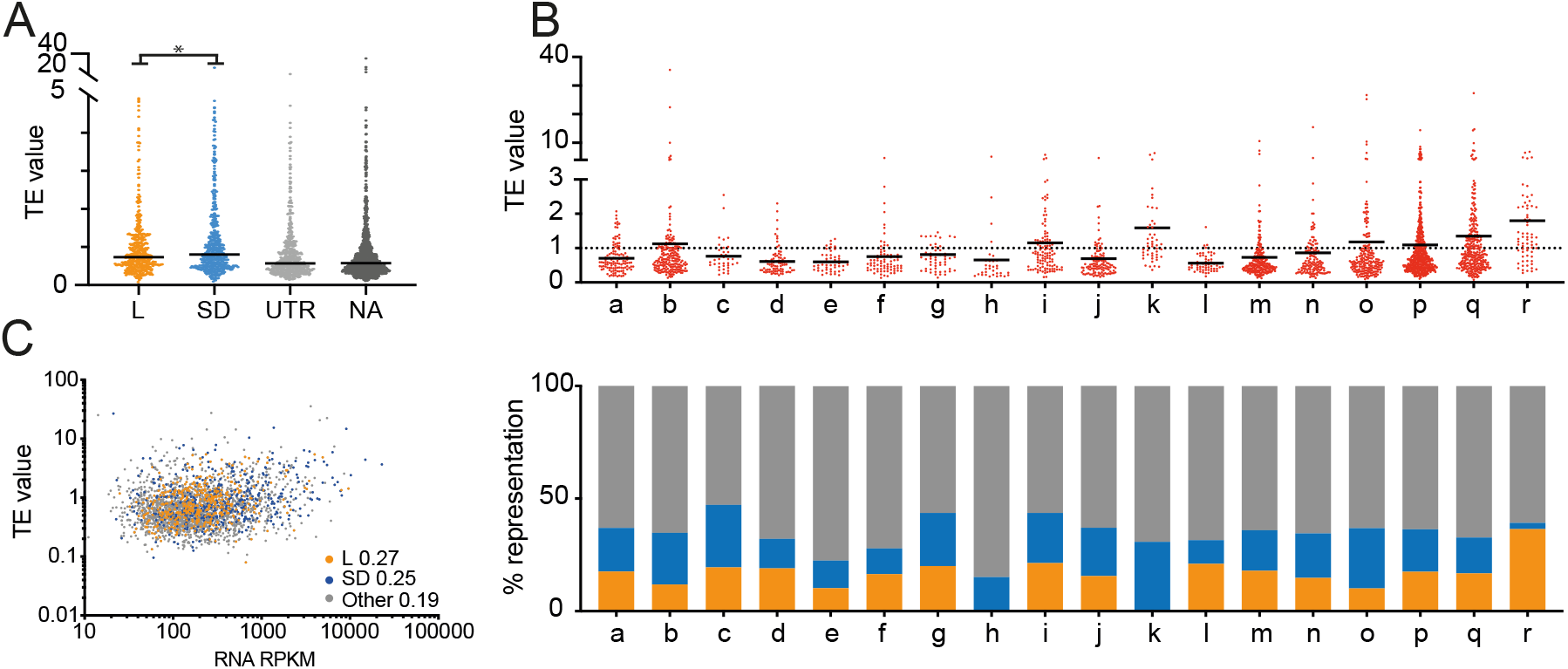
Translation efficiency. (A) Plot of translation efficiency (TE) values distribution of Shine-Dalgarno (SD, blue) and leaderless (L, orange) genes during exponential growth. For reference, genes with a 5’UTR not including a Shine-Dalgarno sequence (UTR, grey) and non-assigned genes (dark grey) are also included. (B) top panel: TE values distribution amongst functional categories (see key below); lower panel: proportion of Shine-Dalgarno (blue) or leaderless (orange) genes representing each functional category (all other genes, including those with a 5’ UTR that lacks a Shine-Dalgarno sequence are shown in grey). With the exception of toxin-antitoxin systems (<u>r</u>) in which the TE values are dominated by a small number of antitoxins with high TEs, in general under conditions of exponential growth, the translational efficiency is highest for genes involved in primary metabolic activities (in which Shine-Dalgarno sequences are dominantly found). Key to functional categories in (B): <u>a</u> Degradation; <u>b</u> Energy metabolism; <u>c</u> Central intermediary metabolism; <u>d</u> Amino acid biosynthesis; <u>e</u> Purines, pyrimidines, nucleosides and nucleotides; <u>f</u> Biosynthesis of cofactors, prosthetic groups and carriers; <u>g</u> Lipid biosynthesis; <u>h</u> Polyketide and non-ribosomal peptide synthesis; <u>i</u> Broad regulatory functions; <u>j</u> Synthesis and modification of macromolecules; <u>k</u> Ribosomal proteins; <u>l</u> Degradation of macromolecules; <u>m</u> Cell envelope; <u>n</u> Cell processes; <u>o</u> Other; <u>p</u> Conserved hypotheticals; <u>q</u> Unknowns; <u>r</u> Toxin-antitoxin systems. (C) Plots of mRNA vs. TE with Spearman coefficients included.

### Metagene analysis reveals differences in ribosome densities at the start codon between leaderless and Shine-Dalgarno transcripts

As leaderless transcripts lack the ribosomal recognition sequences for translation initiation, we hypothesized that maps of ribosome occupancy for genes aligned at their start codon will differ between leaderless and Shine-Dalgarno genes. In order to test this hypothesis, we used ribosome densities assigned to the 3’ end of reads to perform metagene analysis and generate maps of ribosome occupancy. 3’ assignment of reads allowed us to identify a consensus distance of 14 nt from the 3’ end of mapped reads and the start codon offset to the P-site in our dataset (see Methods, Supplementary figure S1), which provides a reliable estimate of the distance from the P-site codon to the 3’ boundary of the mycobacterial ribosome. Visualisation of the metagene profiles obtained for leaderless and Shine-Dalgarno genes revealed striking differences in the ribosome densities around the start codons (Figure 3A). The metagene analysis of Shine-Dalgarno genes shows a peak at the start codon, indicating a high ribosome density in that region, whereas the ribosome density at the 5’ end of leaderless genes increases gradually with distance from the start codon. We propose that the higher density of ribosomes near the start codon of Shine-Dalgarno genes is due to a slower initiation associated with binding of the anti-Shine-Dalgarno and Shine-Dalgarno sequences during positioning of the start codon in the P-site, in contrast to the 70S ribosome binding to leaderless transcripts and initiation proceeding more rapidly to elongation.

To address the question of whether this pausing near the start codon is a global phenomenon that also happens in other bacteria, we analysed previously published Ribo-seq datasets from *M. smegmatis* (Shell et al., 2015) and *E. coli* (Latif et al., 2015) and looked for similarities with our data. Like *M. tuberculosis, M. smegmatis* has a large leaderless transcriptome and appears to be able to translate leaderless transcripts with high efficiency (Shell et al., 2015). In contrast, under optimal growth conditions, *E. coli* expresses only 14 leaderless genes (Thomason et al., 2015). The metagene analysis of the *M. smegmatis* data follows the same pattern as that of our *M. tuberculosis* dataset, suggesting that different species of mycobacteria may share common mechanisms for translating leaderless mRNAs (Figure 3B). However, the same pattern was not seen in *E. coli*, where the metagene analysis of leaderless genes does not appear to indicate an efficient transition between initiation and elongation or indeed reliable translation along the length of the leaderless transcript (Figure 3C).

**Figure 3.**
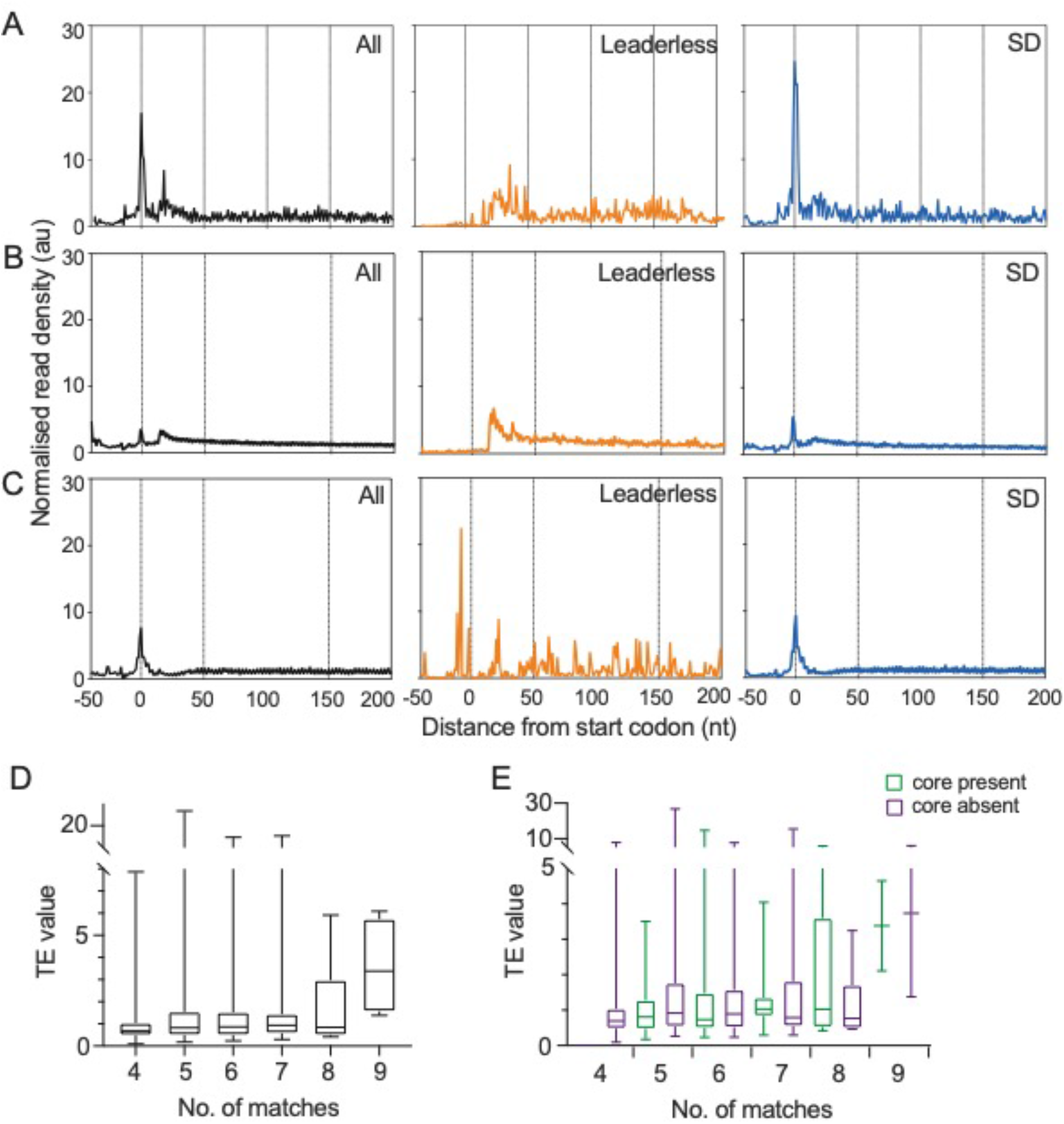
Metagene analysis of 3’ assigned genes aligned at the start codon. (A) *M. tuberculosis* (B) *M. smegmatis* (C) *E. coli* with all genes shown in black, leaderless in orange and Shine-Dalgarno genes in blue. (D) influence of the degree of complementarity to the anti-Shine-Dalgarno on translation efficiency (TE). (E) Influence of the presence of the GGAGG core and degree of complementarity of the anti-Shine-Dalgarno sequence on TE.

TE has been shown to be determined by both codon bias and mRNA folding energy (Tuller et al., 2010). To rule out the possibility that our results were unduly influenced by codon bias, we analysed the codon composition of the 20 codons downstream of the start codon and compared leaderless and Shine-Dalgarno sequences. In general, there were very few significant differences between the types of transcript, although it was interesting to note that it was often the most and least abundant codons genome-wide that were significantly enriched between types of transcript (e.g. the codons CUG and UUA are the most and least used leucine codons in mycobacteria (Andersson and Sharp, 1996), and both were significantly over-represented (Fisher’s exact test P <0.0379; two-sided) in leaderless compared to Shine-Dalgarno transcripts in our dataset (Supplementary Figure S4).

We also investigated the effect of the strength of the Shine-Dalgarno sequence on TE. Locations of Shine-Dalgarno sequences were found by searching for the canonical *M. tuberculosis* Shine-Dalgarno sequence (AGAAAGGAGG; complementary to the anti-Shine-Dalgarno sequence in the 16S ribosomal RNA of *M. tuberculosis* (Kempsell et al., 1992)) in the 30 nucleotides upstream of the start codon. The distance of the Shine-Dalgarno sequence from the start codon and the number of matches between the gene-specific and canonical Shine-Dalgarno sequence were extracted. These metrics were used to assess the relationship between TE and Shine-Dalgarno sequence strength. With the exception of Shine-Dalgarno sequences that are near-perfect matches to the 16S rRNA anti-Shine-Dalgarno sequence (9 complementary base pairs or more, linear regression TE vs. number of matches, P = 0.012) there was no evidence that higher complementarity to the anti-Shine-Dalgarno resulted in higher translation efficiency genome-wide (Figure 3D). It is not possible to say from our data whether for individual genes there is any Shine-Dalgarno complementarity effect, as observed by Park et al. (Park et al., 2011) as we did not specifically test this with mutagenesis; however, there is no apparent correlation between Shine-Dalgarno strength and TE across the genome. The conservation of a core GGAGG motif within the Shine-Dalgarno sequence also had no effect on TE (Figure 3E).

### Transcriptional and translational gene regulation during nutrient starvation

The transcriptional response of the *M. tuberculosis* leaderless transcriptome has been previously shown to increase under conditions of nutrient starvation (Cortes et al., 2013) but its overall effect on the proteome remained unclear. To assess the translational response of *M. tuberculosis* to nutrient starvation and further quantify the effect of leaderless translation on the stress translatome, we performed parallel RNA-seq and Ribo-seq on triplicate *M. tuberculosis* cultures that had been nutrient-starved for 24 hours (see Methods, Supplementary tables 3 and 4). RNA-seq analysis of nutrient starved samples confirmed previous transcriptomic studies with mRNAs for 560 genes increasing > 2-fold and 589 genes decreasing > 2-fold (adjusted P < 0.01, Supplementary table 5), with down-regulated genes including genes coding for ribosomal proteins and genes involved in energy metabolism (Betts et al., 2002; Dahl et al., 2003; Cortes et al., 2013). The global response at the level of the translatome was similar, with RPFs for 563 genes increasing > 2-fold and 573 genes decreasing > 2-fold, but the overlap between transcription and translation was moderate. 344 genes were both significantly up-regulated at the level of mRNA and RPFs and 400 genes were significantly down-regulated (Figure 4A), highlighting a potential scope for translational regulation. To further identify translationally regulated genes upon starvation, we looked at the set of genes with only significant changes at the level of RPFs. We identified a set of 28 genes that showed more than 4-fold induction at the level of translation only (Figure 4B). Interestingly, these genes included both leaderless and Shine-Dalgarno transcripts and are mostly related to cholesterol metabolism (Griffin et al., 2011) or are SEC substrates (de Souza et al., 2011; Målen et al., 2010).

We were able to quantify reliable rates of translation for 2,054 genes, of which 388 (19%) were leaderless and 452 (22%) were Shine-Dalgarno genes. Metagene analysis of read densities aligned at the start codon during nutrient starvation revealed similar patterns of ribosomal recruitment as the ones described for exponential growth, with Shine-Dalgarno genes showing higher ribosome occupancy near the start codon and ribosome occupancy for leaderless genes increasing gradually from the start codon (Supplementary figure 4). Of the mRNA transcripts that were more than 10-fold up-regulated in our previous study (Cortes et al., 2013), 72% were also significantly up-regulated at the level of the translatome in this dataset. Furthermore, the levels of global up-regulation of leaderless genes in the translatome under conditions of nutrient starvation was statistically significant (Fisher’s exact test, P <0.0011) (Figure 4C). To better discriminate if leaderless transcripts are translated with differing efficiencies during nutrient starvation, we calculated TE values as described previously and compared their distribution amongst the set of reliably quantified leaderless and Shine-Dalgarno genes. We found that as for exponential growth, median TE values for Shine-Dalgarno genes were significantly higher than for leaderless genes in the starved translatome (Mann-Whitney test, P =0.0001) (Figure 4D).

As potential mechanisms underlying differential translation of leaderless versus Shine-Dalgarno genes could include quantitative differences in the abundance of components of the translational machinery, we looked for preferential translation of ribosomal proteins, ribosome associated factors as well as translation factors during starvation (Supplementary table 4, Supplementary figure 5). Analysis of the differential expression of ribosome-associated factors during starvation highlighted the significant up-regulation of two genes, Rv1738 and RafS (*DESeq2*, (Love et al., 2014)). The function of the product of gene Rv1738 has not been biochemically determined, but structural studies suggest it may function as a ribosome hibernation promoting factor (HPF) as it bears significant structural homology to the HPF proteins from *E. coli* and *Vibrio cholerae*, including docking of the Rv1738 protein dimer into the groove where HPFs bind the 70S ribosome (Bunker et al., 2015). The up-regulation of Rv1738 in starvation is consistent with its potential role as an HPF and with a role in tolerance of early nutrient starvation. Rv2632c bears some homology to Rv1738, but lacks the ring of positively charged residues likely to be important for ribosomal RNA binding (Bunker et al., 2015). Rv2632c was also up-regulated in starvation (log2 fold-change = 1.59) but the up-regulation fell below the threshold of significance (log2 fold-change = 2). RafS (Rv3241c) was also significantly up-regulated in starvation. RafS is homologous to MSMEG_3935, which contains an S30AE domain and a ribosome-binding domain and has been shown to bind mycobacterial ribosomes under conditions of hypoxic stress (Trauner et al., 2012). Biochemical characterisation of RafS including structural studies of this protein bound to mycobacterial ribosomes indicates a mycobacterial-specific interaction which seems to prevent subunit dissociation and degradation during hibernation without the formation of 100S dimer (Mishra et al., 2018).

**Figure 4.**
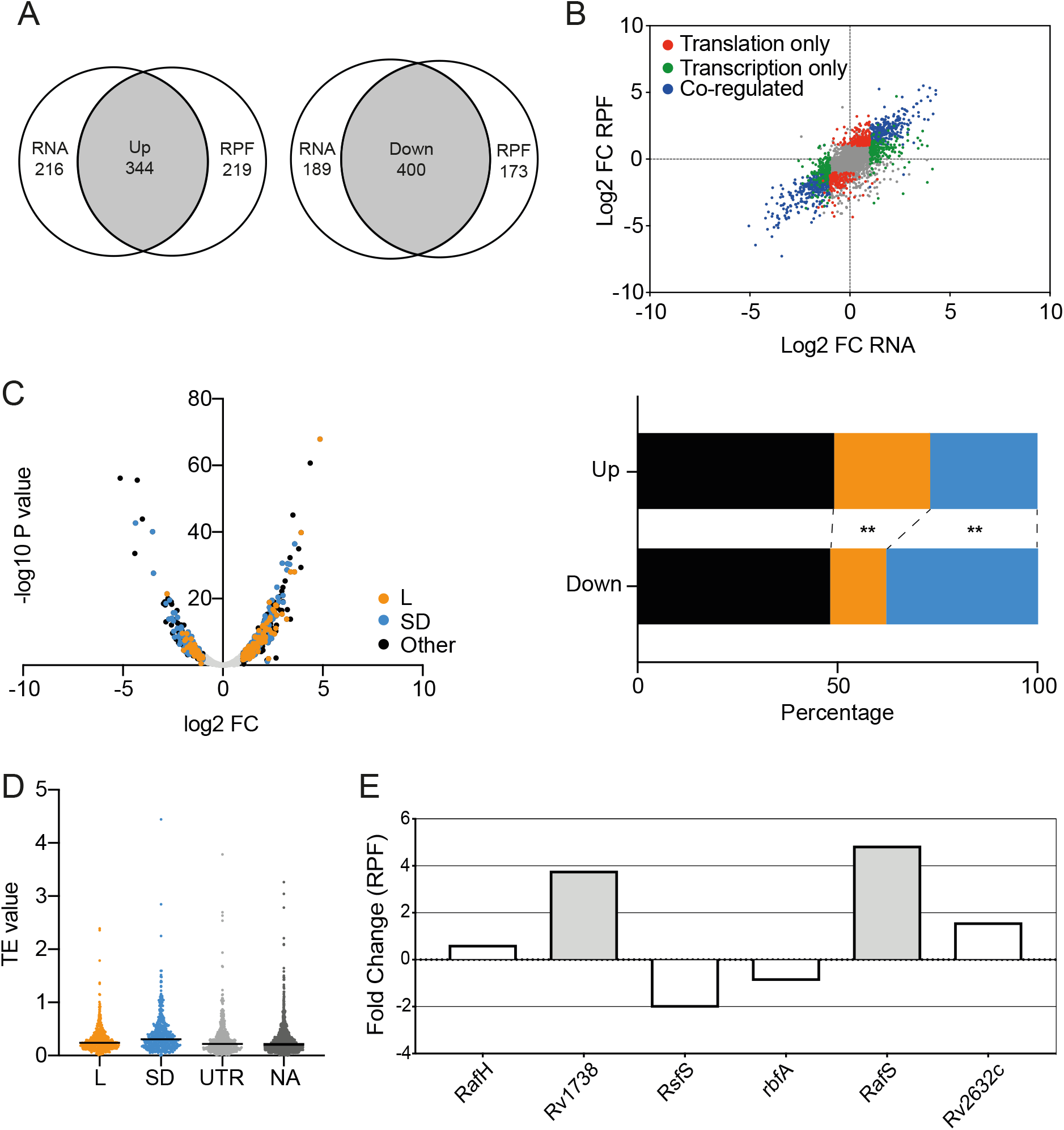
Differential expression in response to starvation. (A) Venn diagrams showing the up- and down-regulation of gene expression at the level of RNA and RPFs. (B) Scatterplot showing the changes in the transcriptome (RNA) and translatome (RPF) during starvation. In blue, co-regulated genes; in red, genes regulated at the level of the translatome only; in green, genes regulated at the level of the transcriptome only. (C) Volcano plot and contingency plots showing the proportion of genes in each UTR category that were up- or down-regulated in starvation. L, leaderless; SD, Shine-Dalgarno. Asterisks denote significant differences (Fisher’s exact test, P = 0.0011) between proportions (D) Distribution of TE values across UTR categories did not change much in starvation compared to control. L, leaderless; SD, Shine-Dalgarno. (E) Changes in the expression of known and putative ribosome-associated factors indicated that Rv1738 and RafS were significantly up-regulated during starvation.

Our data clearly show that translation of leaderless transcripts increases during nutrient starvation, but there was no evidence that this is due to a difference in the TE between different types of transcripts under different conditions.

## Discussion

In this manuscript we present ribosome profiling data from *M. tuberculosis* H37Rv under conditions of exponential growth and nutrient starvation *in vitro*. Our data, which were collected in triplicate, show a high level of reproducibility and appear to represent genuine translation events. There is good agreement with the previously published response of *M. tuberculosis* to nutrient starvation: of the most highly up-regulated mRNA transcripts in nutrient starvation (10-fold or more up-regulation) previously reported (Cortes et al. 2013), 72% were also up-regulated at the level of the translatome. Our data shed light on multiple aspects of translation in mycobacteria, from the role of the Shine-Dalgarno sequence in translation efficiency to mechanisms of initiation.

Our data also provides biochemical evidence for a ~14 nt spacing between the P-site and the 3’ boundary of the mycobacterial ribosome, which is in agreement with estimates obtained in the attenuated *M. tuberculosis* mc^2^7000 strain (Smith et al., 2019) and the distance found in *E. coli* (Woolstenhulme et al., 2015). This distance was obtained by 3’ assignment of RPF reads (Woolstenhulme et al., 2015) which has been shown to be the best method for determining the position of the ribosome on the mRNA. The need for 3’ assignment arises because unlike ribosome profiling libraries from *Saccharomyces cerevisiase*, which show strong three nucleotide periodicity as the ribosome translocates codon by codon along the mRNA, bacterial Ribo-seq libraries show a broad distribution in RPF read lengths. However, when reads are aligned by the 3’ boundary, the start codon peak is reduced to 1-2 nt wide and the position of the ribosome can be accurately inferred (Balakrishnan et al., 2014; Nakahigashi et al., 2014; Woolstenhulme et al., 2015). We therefore used 3’ assignment prior to metagene analysis of our data (and the previously published datasets we analysed from *M. smegmatis* and *E. coli* (Latif et al., 2015; Shell et al., 2015)) to obtain the position of the ribosome.

In our study we used chloramphenicol to arrest translation. Chloramphenicol arrests elongation but has little effect on initiation, potentially leading to an accumulation of ribosome density at the 5’ end of CDSs (Mohammad et al., 2019). To rule out the possibility that our data were affected by chloramphenicol artefacts, we repeated the DESeq2 analysis of differentially translated genes during nutrient starvation to account for reads across the first 50 codons (Supplementary table 6). This revealed little difference in the results, suggesting that in this case any bias towards the 5’ ends of genes has little influence on our analysis.

The role of translational reprogramming as a mechanism for shaping the proteome during stress to guarantee bacterial survival has been widely described in bacteria (reviewed in (Starosta et al., 2014)). In the context of *M. tuberculosis*, the robust translation of leaderless mRNA transcripts during exponential growth and their up-regulation under stress conditions is likely to play a role in cell survival. Our data point towards mycobacterial ribosomes being somewhat promiscuous in terms of the types of mRNA transcript they are able to translate with both leaderless and Shine-Dalgarno transcripts being efficiently translated. Metagene analysis revealed differences in the initiation mechanisms of translation of these two types of transcript, with Shine-Dalgarno transcripts showing ribosomal pausing at the start codon – presumably representing assembly of the preinitiation complex and Shine-Dalgarno to anti-Shine-Dalgarno binding to stabilise the complex. Whereas initiation of leaderless transcripts lacks this pause, consistent with a model in which preassembled 70S monosomes directly bind the mRNA and become quickly elongation-competent.

It is interesting that the putative ribosome stabilisation factors RafS and Rv1738 were significantly up-regulated during nutrient starvation. This observation is consistent with previous studies that have proposed roles for these proteins in 70S ribosome stabilisation and tolerance of early nutrient starvation and hypoxic stress (Bunker et al., 2015; Trauner et al., 2012). The stabilisation of 70S monosomes is relevant to the increase in leaderless translation under conditions of nutrient starvation because the assembled 70S ribosome has been proposed as the species that initiates leaderless translation. This is further supported by our metagene analysis, which indicates a smoother and more rapid initiation for leaderless translation compared to Shine-Dalgarno translation, which exhibits a characteristic pause at the start codon that we attribute to assembly of the pre-initiation complex involving binding of the Shine-Dalgarno-anti-Shine-Dalgarno sequences, 50S subunit joining and GTP hydrolysis to render the ribosome elongation-competent. Similar mechanisms for initiation seem to occur in both exponentially growing and nutrient starved cultures; however, the proportion of leaderless-translated mRNA increased during nutrient starvation. This observation suggests that there are not distinct mechanisms of translation occurring during different growth conditions, but rather that factors that stabilise intact ribosomes may play a role in ensuring continued robust translation during conditions of stress

The influence of the strength of the Shine-Dalgarno sequence on translation efficiency (TE) is a complex issue. Previous studies have reported little or no correlation between ribosome occupancy and the strength of the Shine-Dalgarno motif (Schrader et al., 2014; Li et al., 2014; Li, 2015; Del Campo et al., 2015) when looking at genome-wide effects. However, at the level of individual genes, mutation of the Shine-Dalgarno sequence to decrease binding of the mRNA to the ribosome has been shown to reduce protein expression (Park et al., 2011). How can these results be reconciled? A key study by Buskirk and co-workers dissected out the influence of A-rich sequences in 5’ UTRs from the impact of Shine-Dalgarno strength (Saito et al., 2020). In our study, we found that closer complementarity to the anti-Shine-Dalgarno sequence results in higher translation efficiency only for the highest number of matches; there was little difference in the TEs of genes with imperfect Shine-Dalgarno sequences (according to complementarity with the anti-Shine-Dalgarno sequence (Kempsell et al., 1992)). The four genes with high Watson-Crick complementarity to the anti-Shine-Dalgarno had statistically significant higher TEs than genes with fewer matches. These genes include two conserved hypothetical proteins (Rv0060 and Rv1813c), a probable transcriptional regulatory protein (Rv2324) and the 6 kDa early secretory antigenic target EsxA (Rv3875). As TE is a measure of RPF/mRNA and cannot account for protein degradation, the proteins with high TEs are not necessarily those most abundant in the cell. However, in terms of the mechanism of their translation it is interesting to ask what the transcripts encoding these proteins have in common, and our data indicate that high complementarity of the Shine-Dalgarno sequence is one such feature.

To date, the development of novel therapeutics against *M. tuberculosis* has been slow, hindered in part by the difficulties of studying a slow-growing category 3 pathogen and by the challenges of genetic manipulation of this organism. The data obtained from our experiments provides a rich resource for studying translation and its regulation in *M. tuberculosis* which will be useful to other researchers.

## Materials and Methods

All materials and reagents were of analytical grade and obtained from Sigma-Aldrich (Merck, Darmstadt, Germany) unless otherwise specified. All buffers were prepared using water purified to a resistance of 18.2 MΩ cm^-1^ and filtered through a 0.2 μm membrane (Millipore, Billerica, MA, USA) before use.

### Culture conditions

*M. tuberculosis* H37Rv was grown in Middlebrook 7H9 medium (BD Diagnostics) supplemented with 10% Middlebrook albumin-dextrose-catalase (ADC) (BD Diagnostics), 0.2% glycerol and 0.05 % Tween 80 at 37°C until OD_600_ was 0.4–0.6. For starvation experiments, exponentially growing bacteria were pelleted, washed twice in PBS and resuspended in PBS supplemented with 0.025% Tyloxapol, and maintained in culture for a further 24 h. Triplicate cultures of exponentially growing and nutrient starved cells were prepared for parallel RNA and Ribosome profiling experiments. For ribosome profiling experiments, chloramphenicol was added to a final concentration of 100 μg/ml two minutes prior to harvesting cells. All bacteria were harvested by centrifugation at 4000 rpm and 4°C in a JS 4.0 swinging bucket rotor (Beckman Coulter, Brea California) and immediately processed as described below.

### Ribosome profiling

Sample preparation for ribosome profiling was carried out according to a modified form of the method of Latif et al.(Latif et al., 2015). All steps were performed on ice or using chilled buffers. *M. tuberculosis* cell pellets were resuspended in lysis buffer (20 mM HEPES, pH 7.4; 200 mM KCl, 1% TritonX-100, 12 mM MgCl_2_, 1 mM CaCl_2_, 1 mg/ml heparin, 100 μg/ml chloramphenicol and 5 U/ml DNase I) and transferred to ribolysing matrix tubes (MPBio). Cell lysis was performed by ribolysing the sample three times for 30 s at power 6.5 (Fast Prep Hybaid FP120HY-230, MPBio). Ribolysing matrix and cell debris were removed by centrifugation at 12,000 rpm (JA-15 1.5 rotor, Beckman Coulter) for 10 minutes.

The ribosome profiling samples were then double filtered through a 0.2 μm membrane (Corning, New York) to ensure sterility before incubation for 45 minutes with 24 μl microccocal nuclease (NEB) and 15 μl DNaseI (Thermo Fisher, Waltham, Massachusetts) to digest mRNAs that are not bound by ribosomes. Nuclease digestion was stopped by addition of 10 μl Superase In (Thermo Fisher) and 5 μl 0.5 M EGTA and cell lysates loaded onto a 34% sucrose cushion. Ribosomes were harvested by centrifugation at 40,000 rpm for 15 h in an SW40Ti rotor (Beckman Coulter). To isolate the ribosome-bound mRNAs or ribosome footprints (RPF), the ribosomal pellet was then resuspended in Qiazol (Qiagen, Hilden, Germany) and small RNA isolated using the miRNeasy mini kit (Qiagen). Isolated RNA was precipitated and resuspended in 10 mM Tris, pH 8.0. Finally, RPF samples were depleted of ribosomal RNA using the RiboZero rRNA removal kit for bacteria (Illumina) and purified using RNeasy MinElute columns (Qiagen). The quantity and quality of the RNA was assessed at each step using the bioanalyser and qubit.

### RNA isolation

*M. tuberculosis* cell pellets were resuspended in RNApro solution (MPBio, Irvine, California) and ribolysed three times for 30 s at power 6.5 (FastPrep Hybaid FP120HY-230, MPBio). Ribolysing matrix and cell debris were removed by centrifugation at 12,000 rpm (JA-15 1.5 rotor, Beckman Coulter) 4°C for 10 minutes. The total RNA sample was then removed from the ribolysing tube and chloroform added. The sample was vortexed for 10 s and incubated at room temperature for 5 minutes before centrifugation at 12,000 rpm (JA-15 1.5 rotor, Beckman Coulter) 4°C for 5 minutes. The aqueous phase was then transferred to a new tube and 1.5 volumes of 100% ethanol added. Samples were incubated at −20°C overnight and spun to pellet the RNA. The pellet was washed with 75% ethanol, air dried and resuspended in nuclease-free water. A further acid-phenol chloroform extraction was performed to improve purity of the RNA before resuspending the final pellet in nuclease-free water. Genomic DNA contamination was assessed by PCR for 16S genomic DNA and, if present, samples were treated with DNase I followed by a further acid-phenol chloroform extraction.

Next, samples were depleted of ribosomal RNA using the RiboZero rRNA removal kit for bacteria (Illumina) and purified using RNeasy MinElute columns (Qiagen). Finally, total RNA samples were then subjected to fragmentation using RNA fragmentation reagents (Thermo Fisher) at 80°C for 20 minutes. The quantity and quality of the samples were assessed at each step using the bioanalyser and qubit.

### Library construction and sequencing

RPF and total RNA samples were converted to cDNA libraries as previously described (Latif et al 2015). Briefly, samples were treated with T4 PNK (NEB) at 37ºC for 30 mins and purified using RNeasy MinElute columns (Qiagen). The NEBnext small RNA library prep kit (NEB) was used according to manufacturer’s guidelines to construct the multiplexed cDNA libraries. Multiplexed sequencing was performed by Cambridge Genomic Services using a NextSeq 500 (Illumina, San Diego, California) with single end reads of 75 bp.

### Genome mapping

For the RPF reads, adaptors were removed from the 3’end of sequencing reads using Cutadapt (Martin, 2011) and only trimmed sequences longer than 20nt were kept for further analysis. Bowtie (Langmead et al., 2009) with default parameters was then used to align the trimmed sequences against a reference database containing *M. tuberculosis* ribosomal RNAs (rRNAs) and stable RNAs sequences downloaded from the Mycobrowser website (Kapopoulou et al., 2011). All aligned reads to rRNA were discarded, and the pool of rRNA-depleted RPF reads were finally aligned to the *M. tuberculosis* H37Rv reference genome (AL123456) using Bowtie and allowing for up to two mismatches in a seed length of 28 nt. Reads not aligning to a unique location were discarded.

For the total RNA sequencing reads, Trimmomatic software (Bolger et al., 2014) was used to remove bases from the start and the end of the reads when its quality was below 20, and trimmed reads were mapped against the *M. tuberculosis* H37Rv reference genome as described above.

### Read densities analyses

To gain more insight into ribosome position, a custom phyton script was used to assign ribosome occupancy to the 3’end of RPF reads as previously described (Balakrishnan et al., 2014; Nakahigashi et al., 2014; Woolstenhulme et al., 2015). This method yields higher resolution into ribosome positioning than the previous center-weighting method commonly applied to ribosome profiling studies in bacteria (Becker et al., 2013; Oh et al., 2011). After 3’ assignment, a custom phyton script was used to perform metagene analysis, which gives information on the distribution of ribosomes along an average message by averaging the read densities across all genes (Becker et al., 2013). Only genes greater than 300 nt and which are well-expressed (having more than 100 reads mapped and a minimum number of 1 read per codon along the gene region) were considered for analysis. This was used to generate plots of average ribosome occupancy for genes aligned at their start and stop codons.

In *E. coli*, 3’assignment defines a strong peak of 1-2 nt width located 15 nt downstream of the first nucleotide of the start codon (Woolstenhulme et al., 2015), which corresponds to the distance from the P-site codon to the 3’ boundary of the *E. coli* ribosome (Hartz et al., 1988; Yusupov et al., 2001). In order to determine the P-site offset in our dataset, we used the 3’ assignment to perform metagene analysis at the stat codon for each read length between 20-40 nt, and then measured the distance between the highest 3’ peak of the start codon and the start codon. As shown in Supplementary Figure S1A, this analysis revealed two populations of RPFs, with footprints between 29-37 nt showing clearly defined initiation peaks, indicative of translating ribosomes, and shorter footprints showing less pronounced and defined initiation peaks. As our experimental protocol did not include a size selection step, based on this analysis, we decided to exclude shorter footprints from downstream analysis as they could represent pre-initiation complexes or other artefacts derived from the experimental procedure. After measuring the distances from the 3’end peaks to the start codon we could establish a default p-site offset of 14 nt for our dataset. Next we used this offset value of 14 nt and the set of RPFs from 29-37 nt to re-run the metagene analysis generating a plot of average ribosome occupancy for genes aligned at their start codon that clearly confirmed a pronounced peak at the first nucleotide of the start codon (Supplementary figure S7).

### Quantification of gene expression

For the RPFs and total RNA reads, Bedtools (Quinlan and Hall, 2010) was used to calculate the read coverage for each genomic feature. Mycobrowser annotation (Release R1, November 2017) was used as the annotation reference of the *M. tuberculosis* genome (Kapopoulou et al., 2011). The overall expression rate for each gene was then calculated as reads per kilobase per million (RPKM) values for both RPFs and total RNA reads as previously described (Becker et al., 2013; Mortazavi et al., 2008). For the definition of 5’UTR regions, available data for transcriptional start sites in *M. tuberculosis* (Cortes et al., 2013) was integrated with the Mycobrowser annotation. Reliably quantified genes were those genes with at least 128 total read counts between the three replicates, ensuring that measurements were reliable and not dictated by sampling error (Ingolia et al., 2009)(Supplementary figures S3 and S5C). Translation efficiencies (TEs) for reliable quantified genes were calculated by dividing the expression rates (RPKMs) of translation by those of transcription.

### Differential expression analysis

For the differential expression analysis of RPFs and total RNA reads under nutrient starvation, genome coverage of reads mapping to genes was used for statistical testing using the DESeq2 R package (Love et al., 2014). Differentially expressed genes were considered when fold changes between exponential growth and starvation were greater or equal to 2-fold and the corresponding adjusted p value was less than 0.01.

### Shine-Dalgarno specificity analysis

The Shine-Dalgarno sequence for each gene was found by searching for the canonical Shine-Dalgarno motif (AGAAAGGAGG) in the 30 nucleotides upstream of the start coding using blast (-task blastn-word_size 4). The number of matches and location of the Shine-Dalgarno sequence was extracted by analysing the high-scoring segment pair output by blast. The association between TE and Shine-Dalgarno strength was characterised by performing linear regression between TE and the number of matches and presence of a cores using the lm function in R.

### Statistical analysis

The Spearman rank coefficient was used to determine correlations. GraphPad Prism version 8 for Mac (GraphPad Software, San Diego, California) was used to calculate all statistical tests as described in the text unless otherwise stated. Non-parametric tests were used to evaluate differences among TE comparisons and any other data that were not normally distributed.

### Data availability

RNA-seq and Ribosome profiling data have been deposited in the ArrayExpress database at EMBL-EBI (www.ebi.ac.uk/arrayexpress) under accession number E-MTAB-8835.

## Author contributions

EBS and TC designed the research. EBS performed the experiments. JEP developed bioinformatic scripts, under the supervision of TGC. EBS, JEP and TC analysed the data. EBS and TC wrote the paper. TGC reviewed the paper. All authors read and approved the final manuscript.

## Acknowledgements

The authors thank and acknowledge the support of Cambridge Genomic Services (Department of Pathology, University of Cambridge) who carried out the library quality control, pooling and sequencing. We thank Brendan Wren, Kristine Arnvig and Sam Willcocks for critical reading of the manuscript. This work was supported by funding from the European Research Council (ERC) under the European Union’s Horizon 2020 research and innovation programme (grant agreement No 637730). TGC is funded by the Medical Research Council UK (Grant no. MR/M01360X/1, MR/N010469/1, MR/R025576/1, and MR/R020973/1) and BBSRC UK (Grant no. BB/R013063/1).

