## Supplementary Files for "Ribosome profiling in *Mycobacterium tuberculosis* reveals robust leaderless translation"

The following data tables are available as Excel files:

Table 1 Mapping statistics for data from exponential cultures
Table 2 Reads per kilobase per million (RPKM) and count data from exponential cultures
Table 3 Mapping statistics for data from nutrient-starved cultures
Table 4 Reads per kilobase per million (RPKM) and count data from nutrient-starved cultures
Table 5 DESeq2 data for RNA-seq and Ribo-seq
Table 6 DESeq2 data for RNA-seq and Ribo-seq excluding the first 50 amino acids.

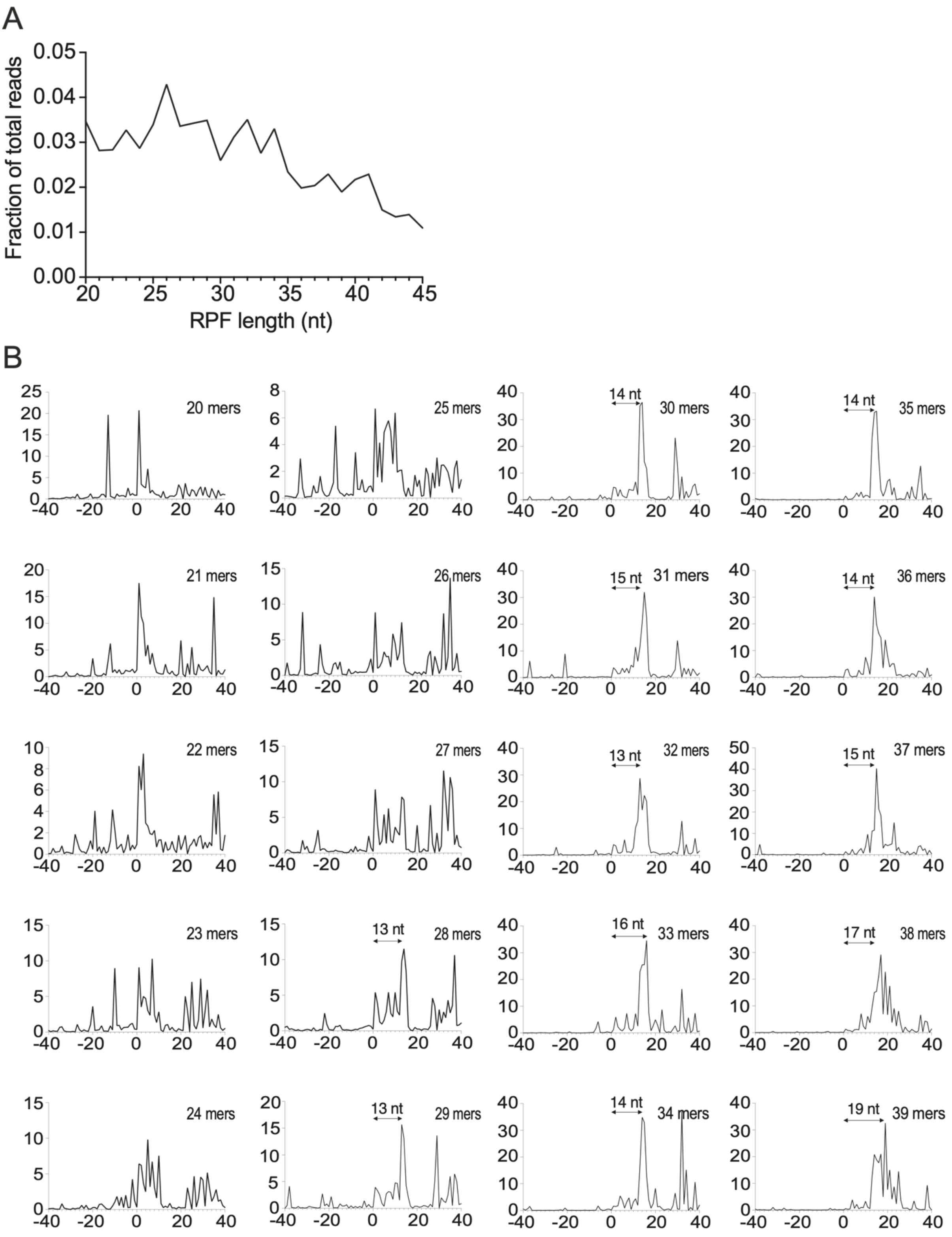


Figure S1
(A) length distribution of RPF reads after adapter trimming
(B) metagene analysis of different RPF readlengths aligned at the start codon (0 nt)

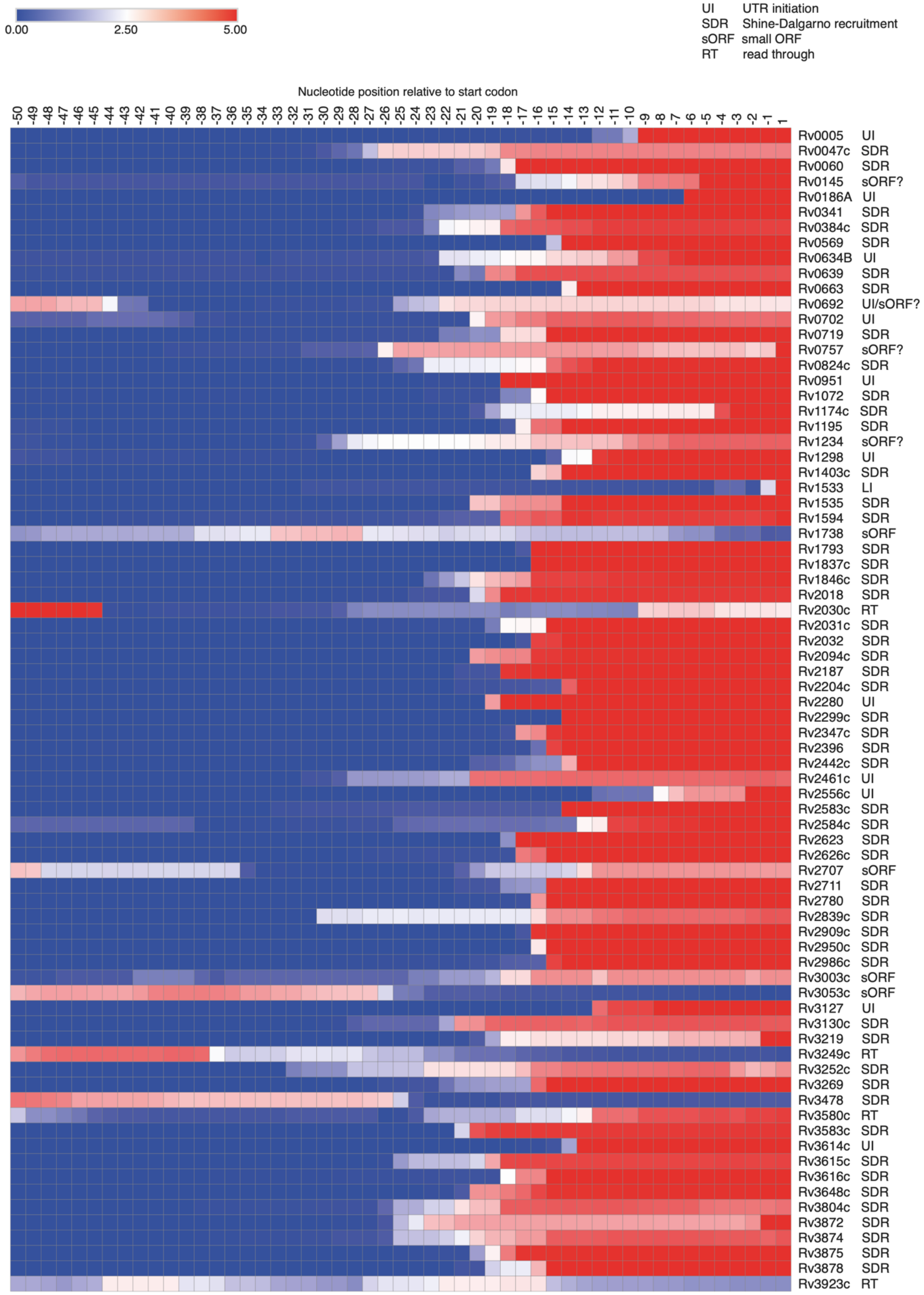


Figure S2
Heatmap of RPF density in the 5' UTRs of the 76 most highly expressed UTRs. Regions of highest density (red) mostly relate to ribosome recruitment, but some genes show read-through from the preceding gene or possible sORFs upstream of the CDS.

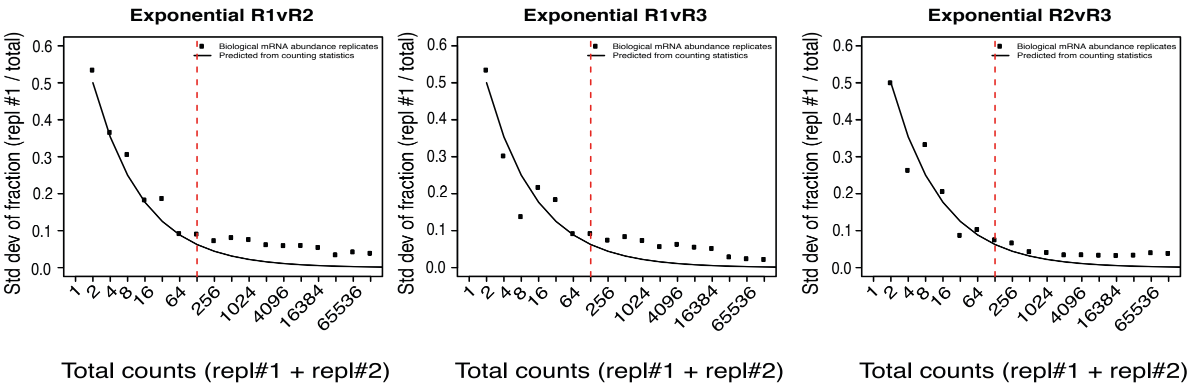


Figure S3 Effects of counting statistics on error in quantification. Pairwise comparisons of the three biological replicates were used to measure reproducibility.

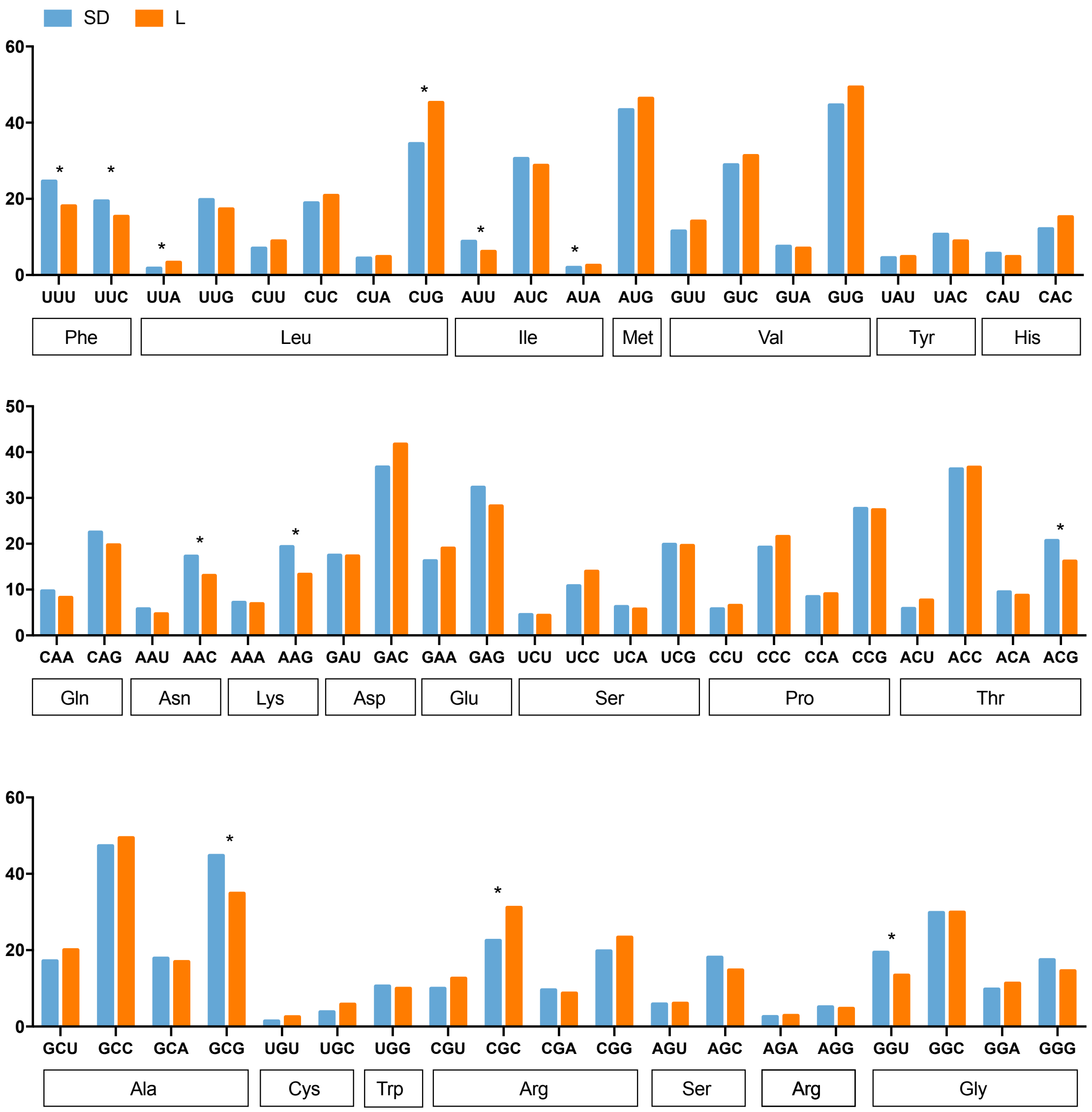


Figure S4 Codon usage analysis

Codon usage in the first 20 codons after the start codon for leaderless (orange) and SD (blue) genes

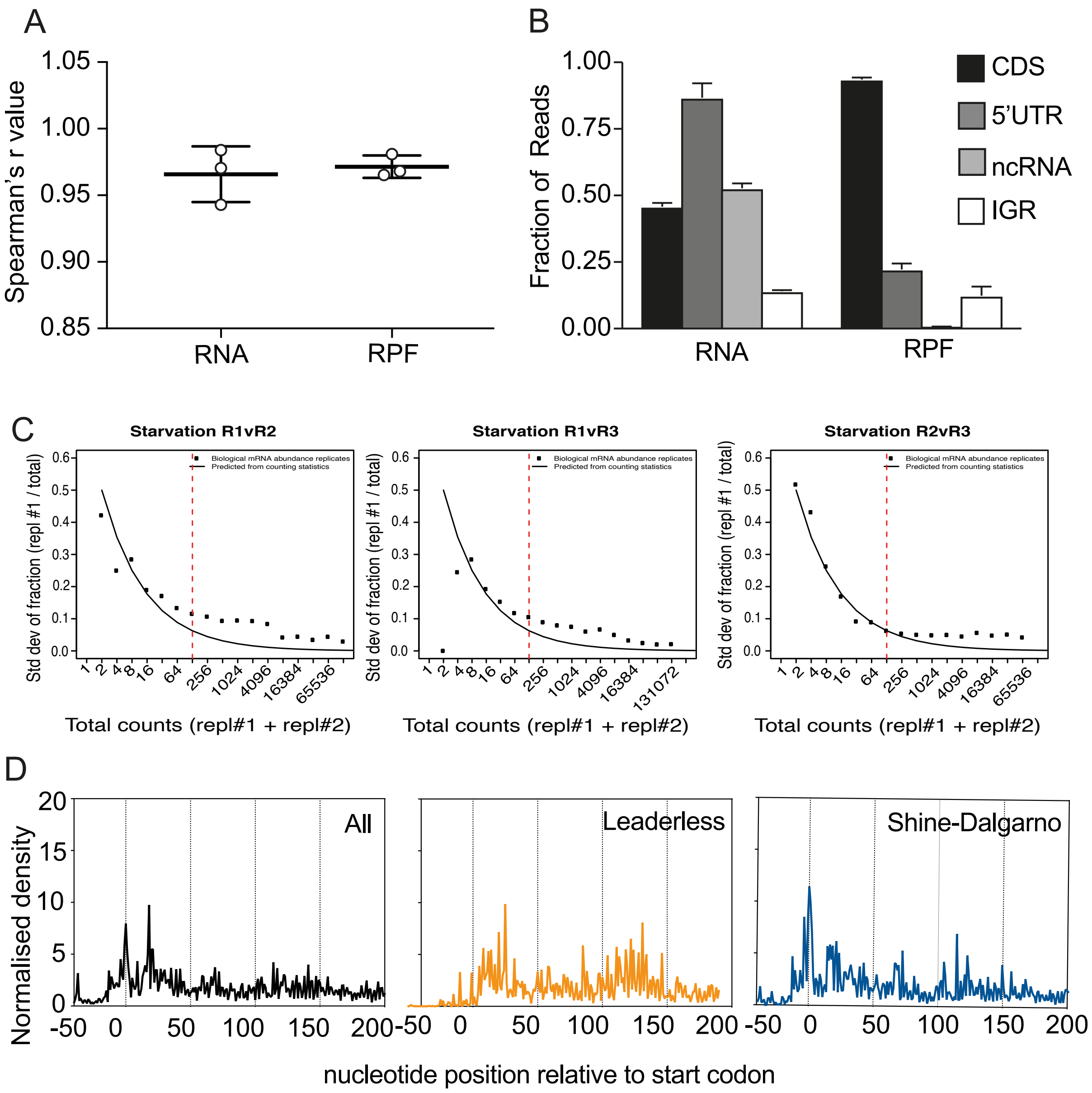


Figure S5 (A) Reproducibility of data obtained for nutrient-starved cultures was similar to that obtained for exponentially growing cultures. (B) Reads mapped to various genomic regions also showed a similar pattern to the exponentially growing cultures. (C) Effects of counting statistics on error in quantification. Pairwise comparisons of the three biological replicates of starvation cultures were used to measure reproducibility. (D) Metagene analysis showed the characteristic peak at the start codon for SD genes (blue) and a smoother initiation pattern for leaderless genes (orange)

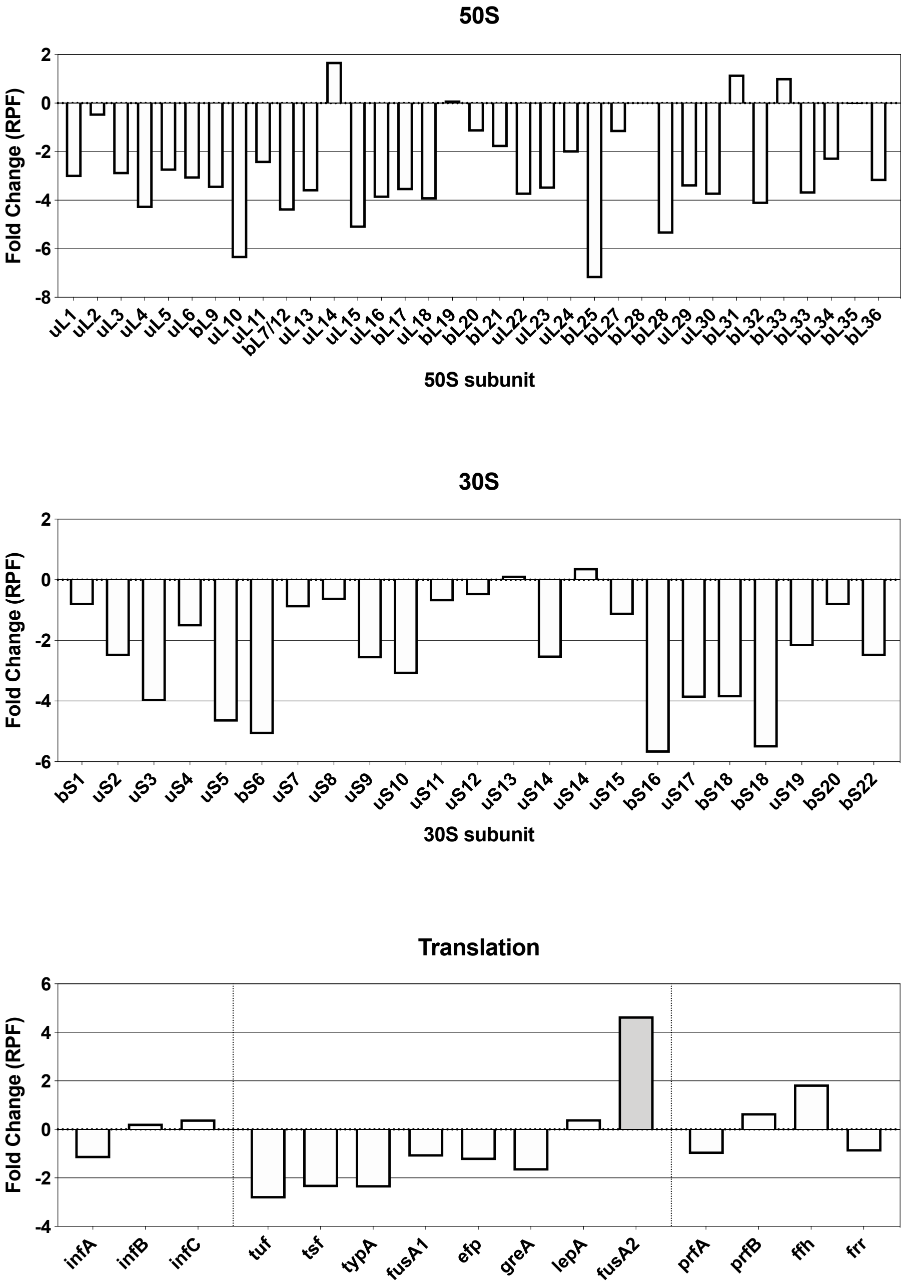


Figure S6 Changes in expression of translation machinery and associated factors during starvation

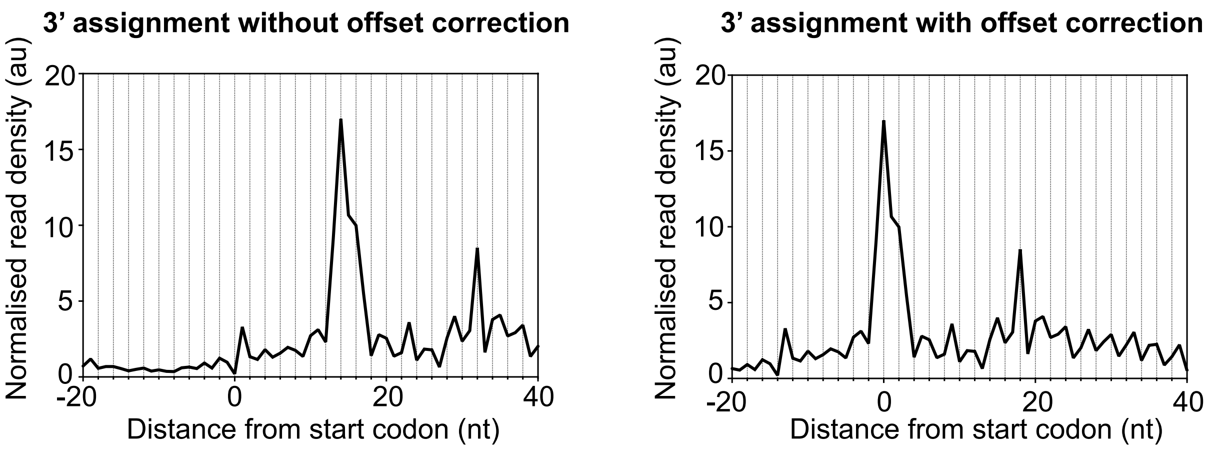

Figure S7 A 14 nt offset correction after 3’ assignment places the start codon in the P-site of the ribosome, indicated by high ribosome density at the start codon as the initiation complex becomes elongation-competent.
